# PXL: a Nucleic Acid–Binding Module of Promyelocytic Leukemia Protein

**DOI:** 10.64898/2026.02.19.706916

**Authors:** Daniel Fairchild, Irina V. Semenova, Dane Geddes-Buehre, Yunfeng Li, Renata Szczepaniak, Sandra K. Weller, Bing Hao, Irina Bezsonova

## Abstract

The promyelocytic leukemia protein (PML) is a stress-response factor that assembles into PML nuclear bodies, dynamic subnuclear compartments involved in tumor suppression and antiviral defense. The most abundant isoform, PML-1, has been linked to transcriptional regulation, genome stability, and antiviral responses, yet the molecular basis of these functions remains unclear. Here, we report that PML-1 contains a unique nucleic acid– binding module, PXL, and determine its three-dimensional structure by X-ray crystallography. Further biochemical, mutational, and cellular analyses, including RNA-seq, demonstrate that this module selectively binds single-stranded G-rich RNA and DNA motifs and modulates the transcriptome. These findings reveal an unexpected molecular function of PML and provide a framework for understanding its roles in nuclear organization and gene regulation.

## INTRODUCTION

The promyelocytic leukemia protein (PML, or TRIM19) is a SUMO E3 ligase (1) and scaffold protein that organizes spherical subnuclear compartments known as PML nuclear bodies (PML-NBs) (2). These bodies play key roles in cellular stress responses. PML forms the outer shell of PML-NBs and recruits numerous SUMOylated client proteins, including Sp100, DAXX, transcription factors, and DNA repair proteins, into the interior (3,4). These dynamic compartments act as hubs that coordinate transcription, DNA repair, antiviral defense, and genome maintenance (4-6). Consistent with these roles, PML-NBs have been observed near sites of DNA damage (7), regions of nascent RNA transcription (8), and viral DNA (6). PML also contributes to the innate antiviral immune response, and many DNA viruses, including herpes simplex viruses, target and disrupt PML-NBs (9-11). Viral genomes of herpesviruses and adenoviruses often localize adjacent to PML-NBs, which have been shown to have an inhibitory effect on viral replication (3,9). Despite these well-established associations, the molecular mechanisms by which PML-NBs execute these diverse functions remain poorly understood.

There are seven primary PML isoforms (1-7) produced by alternative splicing of the *PML* gene (5). They share a common N-terminal RBCC motif, which consists of RING, B-box1, B-box2, and coiled-coil domains (RBCC), but differ in their C-termini (**Figure 1A**). While the RBCC motif is essential for PML-NB formation, the variable C-termini mediate isoform-specific interactions with nuclear components (12). PML-1 is the most abundant isoform (13). The isoform-specific functions of PML-1 include regulating transcription factors, targeting nucleoli, maintaining genomic stability, and responding to DNA viruses (14-16). It is highly expressed in breast cancer, where its levels correlate with poor prognosis (13). ChIP-seq analyses show that, unlike other PML isoforms, PML-1 localizes to promoters of stemness genes, enhancing the stem-like properties of cancer cells (13). These findings suggest that the PML-1 isoform contains a unique module that directs it to specific genomic loci.

**Figure 1:**
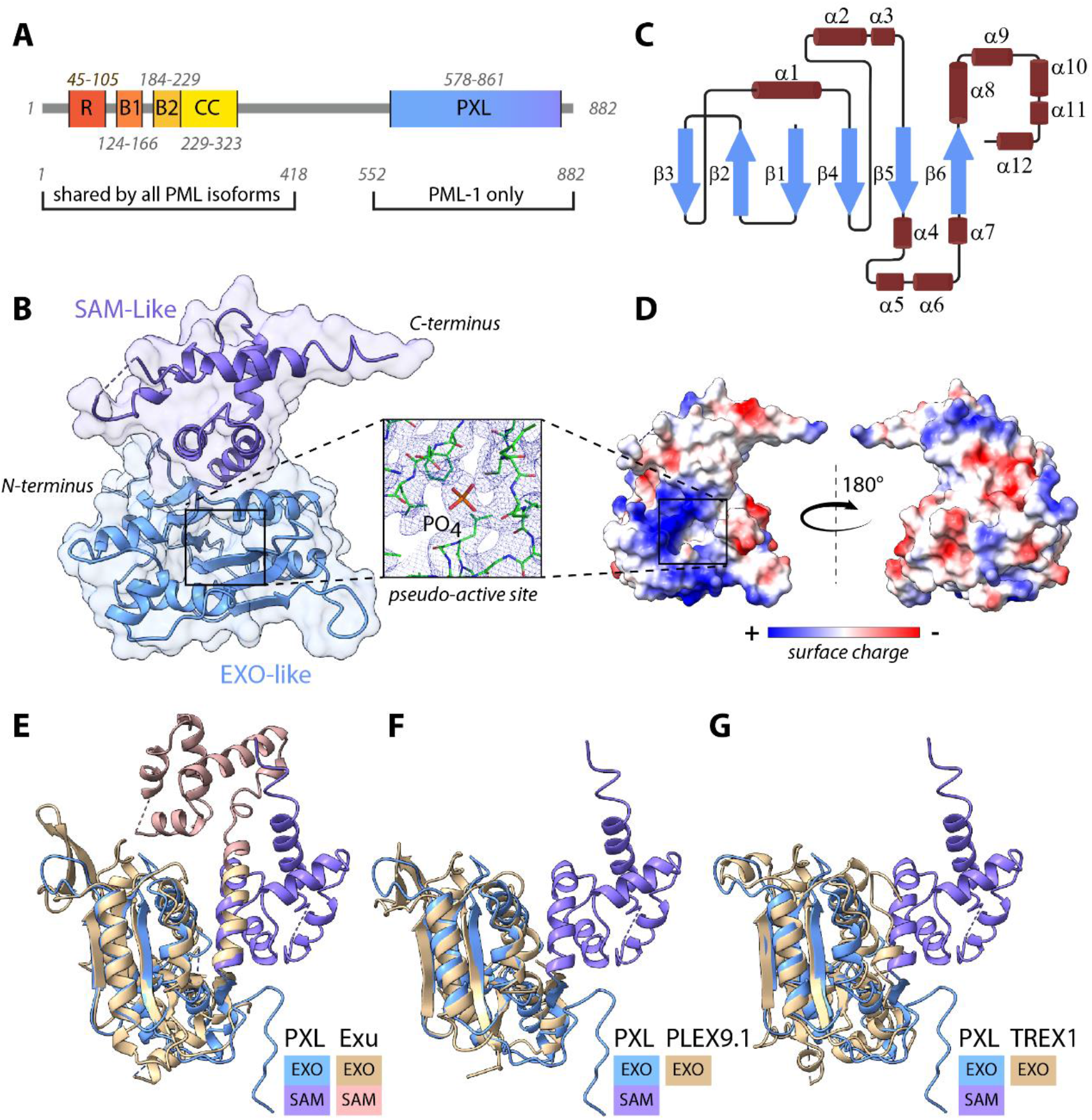
Isoform 1 of PML contains an exonuclease-like domain, PXL. **(A**) Schematic domain organization of PML-1, which consists of N-terminal Ring (R), Bbox1 (B1), Bbox2 (B2), and coiled coil domain shared by all PML isoforms, and isoform 1-specific C-terminal PXL domain. **(B)** Crystal structure of the PXL domain with two subdomains labeled, EXO-like (light blue) and SAM-like (purple). The inset shows the electron density of the pseudo-active site of the EXO sub-domain with the phosphate ion labeled. **(C)** Secondary structure diagram of PXL. **(D)** Surface representation of PXL domain in two orientations, colored by surface charge from positive (blue) to negative (red). An extensive positively charged surface patch (blue) surrounds the pseudo-active site (square). **(E-G)** Structural overlay of PXL and its closest structural homologs found in the Protein Data Bank: **(E)** Exu (Z-score: 19.1, RMSD: 2.7), **(F)** PLEX9.1 (Z-score: 17.9, RMSD: 2.1) and **(G)** TREX1 (Z-score: 15.1, RMSD: 2.7). Structures and sub-domains are color-coded and labeled.

The unique C-terminal sequence of PML-1, encoded by exon 9 of the *PML* gene, includes a putative nuclease domain (14) (**Figure 1A**). This region has been linked to nucleolar targeting, cellular senescence, and maintenance of genomic stability (14,15). To date, nucleic acid binding, or DNase/RNase activity have not been explicitly demonstrated for human PML-1. Here, we show that PML-1 contains a pseudo-nuclease domain, PXL (**P**seudo-e**X**onuclease **L**ike), a nucleic-acid–targeting module unique to isoform 1. Structural and functional analyses, including X-ray crystallography, biochemical, biophysical, and RNA-seq reveal that PXL directly and specifically binds single-stranded G-rich RNA and DNA motifs *in vitro* and influences the cellular transcriptome. Together, these findings establish a direct nucleic acid–binding function for PML-1 and provide a mechanistic framework for its roles in transcriptional regulation, antiviral defense, and genome maintenance.

## MATERIALS and METHODS

### Protein Expression and Purification

The PXL gene sequence was codon-optimized for bacterial expression and chemically synthesized (Genscript). Ligation-independent cloning was used to insert the PXL gene (578-861Δ5) into a pET-based vector containing TEV-cleavable His6 and SUMO tags. The vector was a gift from Scott Gradia (Addgene plasmid #29711). To generate PXL(578-861Δ5)2K1R/3D, R792D, K678D, and K725D mutations were introduced into the PXL(578-861Δ5) sequence using site directed mutagenesis. The PXL(578-861Δ5) construct was transformed into C43 strain of *E. coli* and plated on LB medium with ampicillin. An isolated colony of the transformation was used to inoculate 50 mL of LB broth for an overnight growth at 37 °C. From the overnight growth, 25 mL was used to inoculate 1 L of LB media containing ampicillin and grown at 37 °C to OD_600_ of 0.8 a. u. Protein expression was induced overnight at 18 °C by the addition of 1 mM IPTG. Cell pellets were resuspended in 20 mM NaH_2_PO_4_ pH 8.0, 500 mM NaCl, 5 mM BME, and 15 mM imidazole, then lysed by sonication. The insoluble components of the lysate were removed via centrifugation at 15,000 rpm in a Sorvall RC 6+ centrifuge. The soluble lysates were then purified on a HisPur cobalt resin column and eluted using 20 mM NaH_2_PO_4_ pH 8.0, 500 mM NaCl, 5 mM BME, and 400 mM imidazole. Eluted proteins were dialyzed into 20 mM Tris-HCl pH 7.5, 100 mM NaCl, 5 mM BME and cleaved with TEV protease overnight at 4 °C. After dialysis and cleavage, PXL(578-861Δ5) was further purified on a HiTrap Heparin HP affinity column using linear gradient of NaCl (0.1-1.0 M) in 20 mM Tris-HCl pH 7.5 and 5 mM BME. Eluted fractions were then collected and analyzed by SDS PAGE. Fractions containing purified PXL(578-861Δ5) were pooled and buffer exchanged into 20 mM Tris-HCl pH 7.5, 100 mM NaCl, and 5 mM BME.

The TREX1 expression plasmid was a gift from Cheryl Arrowsmith (Addgene plasmid # 220884). TREX1-6xHis was transformed into the Bl21 (DE3) strain of *E*.*coli* and plated on LB agar containing kanamycin. An isolated colony was selected and used to inoculate 50 mL of LB containing kanamycin for overnight growth at 37 °C. From the overnight, 25 mL were used to inoculate 1 L of LB with kanamycin and was grown at 37 °C until it reached an OD_600_ of 0.8 a.u. Protein expression was then induced overnight at 18 °C by the addition of 1 mM IPTG. Cell pellets were resuspended in lysis buffer (50 mM NaH_2_PO_4_, 300 mM NaCl, 10 mM imidazole, pH 8.0) and lysed by sonication. The insoluble components of the lysate were removed via centrifugation at 15,000 rpm in a Sorvall RC 6+ centrifuge. The soluble lysates were then purified on a HisPur cobalt resin column and eluted using 300 mM imidazole in lysis buffer (pH 8.0). Eluted proteins were then dialyzed into FPLC buffer (20 mM Tris-HCl, 150 mM NaCl, pH 8.0) overnight at 4 °C, and simultaneously, the 6xHis tag was removed with TEV protease. After dialysis and cleavage, TREX1 was further purified via size exclusion chromatography using a HiLoad Superdex 75 (Cytiva) column in FPLC buffer.

### X-Ray Crystallography

Seed crystals of PXL(578-861Δ5) (10 mg/mL) were first obtained in 2 M NaCl, 100 mM Tris-HCl pH 7.0, and 200 mM MgCl_2_ at 16 °C using hanging-drop vapor diffusion and then crushed through shaking with small glass beads using a vortex. Larger PXL(578-861Δ5) (10 mg/mL) crystals were then obtained in 2M NaCl, 100 mM Tris-HCl pH 6.75, and 200 mM MgCl_2_ at 16°C with seeding through hanging-drop vapor diffusion. Crystals were cryoprotected in reservoir solution supplemented with 20% glycerol and flash-cooled in liquid nitrogen. X-ray diffraction data were collected at NSLS-II beamlines 17-ID-1 and 17-ID-2, and processed using Fast DP (17,18) and autoPROC (19). The crystals contain one molecule in the asymmetric unit. The structure of PXL(578-861Δ5) was determined by molecular replacement in Phaser/CCP4 (20,21) using the structure model generated by AlphaFold3 (22) as a starting model. The models were refined by alternating cycles of manual rebuilding in Coot and refinement with REFMAC5/CCP4 (21,23). Data collection and refinement statistics are summarized in Table 1. Ramachandran statistics were calculated using MolProbity(24), and figures generated in Chimera (25) and PyMOL (Schrödinger, LLC).

**Table 1.** Summary of crystallographic analysis.

|  |  |
| --- | --- |
| Data Collection | PXL(578-861Δ5) |
| Wavelength (Å) | 0.9793 |
| Space group | $P3_121$ |
| Cell dimensions (Å) |  |
| *a*, *b*, *c* (Å) | 114.34, 114.34, 102.68 |
| $\alpha$, $\beta$, $\gamma$ (°) | 90.0, 90.0, 120.0 |
| Resolution (Å) | 33.0-3.01 (3.39-3.01) |
| $R_{\text{sym}}$ (%) | 0.46 (3.75) |
| Mean ($\langle I/\sigma I \rangle$) | 6.0 (1.4) |
| Completeness (%) | 93.3 (67.3) |
| CC (1/2) | 0.992 (0.324) |
| Multiplicity | 12.5 (12.2) |
| Refinement |  |
| Resolution (Å) | 21.61-3.01 |
| No. reflections ($\vert F\vert > 0\sigma$) | 9707 |
| $R_{\text{work}}/R_{\text{free}}$ (%) | 16.7/24.1 |
| No. atoms |  |
| Protein | 2,069 |
| Water | 194 |
| Average B-factors (Å<sup>2</sup>) |  |
| Protein | 87.4 |
| Water | 57.4 |
| Wilson B-factors (Å<sup>2</sup>) | 95.2 |
| R.m.s. deviations |  |
| Bond lengths (Å) | 0.01 |
| Bond angles (°) | 1.18 |
| Ramachandran plot (%) |  |
| Favored (%) | 100 |
| Allowed (%) | 0.0 |
| Outliers (%) | 0.0 |

Values in parentheses are for the highest-resolution shell.

### Exonuclease activity assay

The exonuclease activity binding mixtures contained 500 nM of a linearized 6 kB plasmid with staggered ends, and either 200 nM TREX1 or PXL in 20 mM Tris-HCl pH 8.0, 100 mM NaCl, 5 mM BME, and 5 mM MgCl_2_ or 10 mM EDTA. Binding mixtures were incubated for 0, 15, or 30 minutes and reactions were stopped by the addition of loading dye containing SDS. Binding mixtures were run at 100V for 30 minutes in 0.5% agarose gel in TAE buffer and labeled with SYBR Safe DNA gel stain. Gels were imaged using a Syngene G:Box gel imager.

### Electrophoretic mobility shift assay

The EMSA binding mixtures contained 500 nM 5’ or 3’-AlexaFluor488 labeled nucleotide substrates, 0–3 µM PXL(578-861Δ5), 20 mM HEPES pH 7.3, 150 mM NaCl, 1 mM BME, 2.5 mM MgCl_2_, 100 µg/mL BSA, and 6% ficol. Binding mixtures were incubated on ice for 30 minutes and resolved by gel electrophoresis using a native 5% polyacrylamide TBE gel. Gels were visualized with the BioRad ChemiDoc MP imaging system.

5’ and 3’-AF-488 labeled oligonucleotides were chemically synthesized (IDT); 5’ AF-488 RNA sequences: 5’-UGUGUGUGUGUGUGUGUGUG-3’, 5’-UGUGUGUGUG-3’, 5’-ACACACACAC-3’, 5’–AAAAAAAAAA-3’, 5’-CCCCCCCCCC-3’, 5’–UUUUUUUUUU-3’; and 5’AF-488 ssDNA sequences: 5’-TGTGTGTGTG-3’, 5’-ACACACACAC-3’, 5’-AAAAAAAAAA-3’, 5’-GGGGGGGGGG-3’, 5’-CCCCCCCCCC-3’, 5’-TTTTTTTTTT-3’; and 3’ AF-488 dsDNA sequences: 5’-AGCTATGGCGTCGAA-3’ and 3’-TCGATACCGCAGCTT -5’

### Fluorescence Polarization

FP experiments were performed with 100 nM 5’-AlexaFluor488 labeled nucleotide substrates on a SpectraMax iD5 fluorescence plate reader. Nucleotide substrates were incubated with PXL (578-861Δ5) in 20 mM Tris-HCl pH 7.5, 1 mM BME, 1 mM EDTA and 150 mM NaCl. The total reaction volume was 15 µL. Duplicates of each titration point were prepared and measured with 485 nm and 535 nm as the respective excitation and emission wavelengths. The data were fit using nonlinear regression on the Prism 10 software (GraphPad).

### Cell Lines

PML^-/-^ U2OS cells (26) were grown in McCoy’s 5A modified medium supplemented with 10% fetal bovine serum (FBS) and seeded into a six-well plate at approximately 3*10_4_ cells per well. On the following day, cells were washed twice with PBS and media was replaced with Opti-MEM reduced serum media. Cells were transfected with plasmids containing pUC119 carrier DNA and either YFP-WT-PML-1 or YFP-PML-1(1-600). Transfections were performed using Lipofectamine 2000 according to the manufacturer’s instructions. Transfections were carried out over 24 hours upon which the transfection mixture was removed, and cells were washed 3 times with PBS and grown in McCoy’s 5A modified medium supplemented with 10% FBS for 24 hours. Cells transfected with YFP-WT-PML-1 or YFP-PML-1(1-600) were washed 2 times with PBS and suspended with .05% trypsin. Trypsinization was halted by the addition of growth media supplemented with 10% FBS. Suspended cells were sorted using the FACSymphony S6 cell sorter (BD Biosciences). Single cells positive for YFP signal were plated in a 96-well plate and expanded in growth media supplemented with 10% FBS and 100 ug/mL G418.

### Bulk RNAseq

PML^-/-^ U2OS, WT U2OS, PML_-/-_ U2OS expressing YFP-WT-PML-1 or YFP-PML-1(1-600), were cultured in 100 mm dishes to 90% confluency. Cells were subsequently washed 2X with PBS, resuspended using 0.05% trypsin. Following resuspension, PML^-/-^ U2OS expressing YFP-WT-PML-1 or YFP-PML-1(1-600) cells were sorted by YFP fluorescence using the FACSymphony S6 cell sorter. Cells with YFP signal detected were collected separately and utilized for bulk RNAseq. For all cell lines, 100,000 cells were pelleted at 100 G and washed with PBS. Following washing, cells were pelleted again at 100 G and resuspended in DNA/RNA Shield_TM_ (ZYMO Research) until shipment to PlasmidSaurus (plasmidsaurus.com) for RNAseq sample preparation and data collection. Three biological replicates were used for each cell line in RNAseq.

## RESULTS

### Isoform 1 of PML contains an exonuclease-like domain

Protein sequence analysis of PML isoforms (27) in combination with AlphaFold (22,28) predict a structured domain within the C-terminal region of PML-1 (**Figure 1A** and **Supplementary Figure S1**). This region has been previously predicted to contain a nuclease-like domain (14,15).

To further characterize the putative nuclease domain, we solved its structure by X-ray crystallography (**Figures 1B-D**). The crystallized PML-1 fragment included residues 578-861 with a short deletion (645-FFSIY-649), which significantly improved the domain’s solubility. The structure refined to 3 Å resolution with R_free_ of 24.1% and an R factor of 16.7 % (**Table 1**), revealed two subdomains: an α/β fold resembling 3’-5’ DEDD family exonucleases (29) (EXO-like), and a helical bundle similar to Sterile Alpha Motif (SAM-like) (SCOP 47768). The domain contains a central six-stranded β-sheet (β3-β2-β1-β4-β5-β6), with both parallel and antiparallel strands sandwiched between helical elements α1, α2, α3, and α4, α5, α7, α8 on each side, forming a typical EXO-like domain (**Figure 1C**). Immediately following the β6 strand, a 22 Å-long helix, α8, extends into the SAM-like region, where it contributes to a five-helix bundle with α9-α12 helices (**Figure 1C**). Electrostatic surface analysis revealed an extensive positively charged patch located on one face of the domain (**Figure 1D**), consistent with the nucleic acid-binding properties of nuclease-like folds.

Structural homologues of the domain were identified using DALI (30), and revealed significant similarity to 3’-5’ DEDD family nucleases, including exonuclease domains of several DNA polymerases. The top hits were *Drosophila* pseudo-nuclease Exu^19^ (PDB:5L80, 11% sequence homology), zebra fish 3’-5’ exonuclease Plex9.1 (PDB:9MRC, 15% sequence homology) and human 3’-5’ exonuclease TREX1 (31) (PDB:7TQP, 10% sequence homology) (**Figure 1E-G**). Among structural homologues, only the pseudo-nuclease Exu has a similar EXO/SAM arrangement, although with a different relative orientation of the subdomains (**Supplementary Figure S2**). In contrast, the SAM subdomain has not been found in any other nucleases. Therefore, we designate this PML-1 domain **P**seudo-e**X**onuclease-**L**ike domain, or PXL (pixel).

### PXL lacks nuclease activity

Structural similarity between PXL and exonucleases raises the possibility that PXL may function as an exonuclease. Canonical 3’-5’ exonucleases catalyze processive hydrolysis of nucleic acids from the 3’ end via a two-metal ion-dependent mechanism (29). DEDD exonucleases share a conserved catalytic core with a β1-β2-β3-α1-β4-α2-β5-#x03B1;3 fold and an active site formed by four conserved acidic residues (DEDD), often followed by a basic residue (H or Y), which coordinate magnesium ions and mediate hydrolysis (29).

Despite this structural similarity, sequence analysis argues against a catalytic role for PXL. Multiple sequence alignment of PXL with a pseudo-nuclease Exu and catalytically active exonucleases TREX1 and Plex9.1 reveals that the canonical DEDD residues are not conserved in PXL and Exu but are preserved in active exonucleases (**Figure 2A**). Only one of the four DEDD residues, D614, is retained in PXL (**Figure 2A-B**). The absence of conserved metal-coordinating residues indicates that the PXL domain of PML-1 is unlikely to function as an exonuclease.

**Figure 2:**
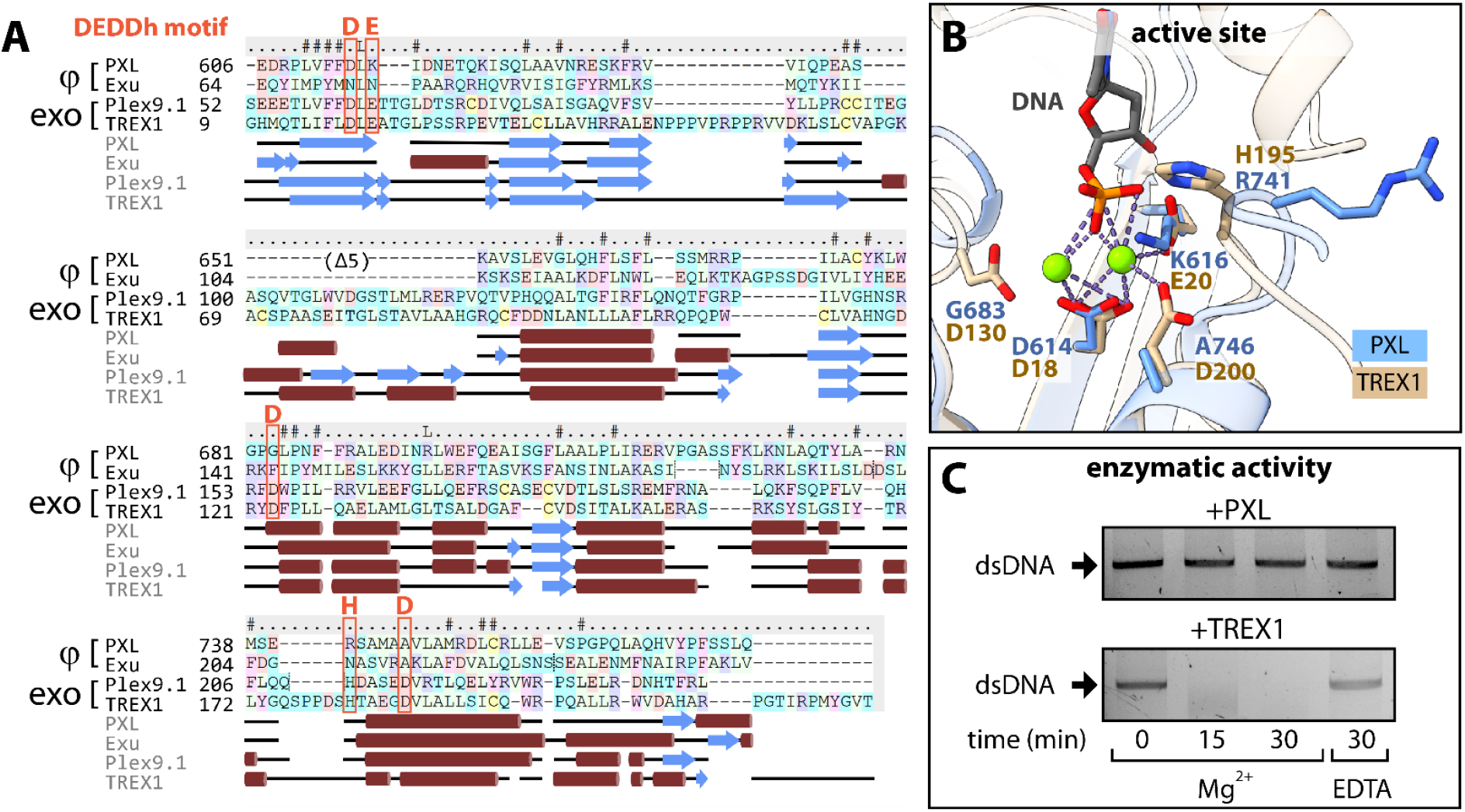
PXL is not an active exonuclease. **(A)** Multiple sequence alignment of the human PML-1 PXL domain, the *drosophila* pseudo-nuclease Exu, zebrafish PML-like exon 9 (Plex9.1), and human TREX1. Predicted secondary structural elements are shown underneath the multiple sequence alignment, alpha helices are depicted as red cylinders, and beta strands as blue arrows. The conserved DEDD nuclease active site is outlined in red. **(B)** Overlay of TREX1 (tan) and PXL (blue) active/pseudo active site. TREX1 catalytic residues and PXL aligned residues are represented as sticks. TREX1 catalytic residues coordinate magnesium ions (green) to facilitate nucleic acid (gray) cleavage. **(C)** Nuclease activity assay showing TREX1 cleaving dsDNA in a metal dependent manner over time. PXL does not show exonuclease activity.

To experimentally confirm that PXL lacks exonuclease activity, we conducted an *in vitro* activity assay using TREX1 as a positive control (**Figure 2C**). As expected, TREX1 was able to completely degrade the dsDNA substrate in ∼30 minutes and its activity was dependent on the presence of magnesium. Under the same experimental conditions, PXL demonstrated no detectable nuclease activity, thus showing that unlike TREX1, PXL does not facilitate nucleic acid cleavage through coordination of magnesium ions.

Consistent with the sequence analysis, the crystal structure of PXL shows no evidence of magnesium ions in the active site despite the presence of magnesium salt in the crystallization solution. Instead, residual electron density is observed buried deep in the pseudo-active site between β1 and a loop connecting the β4 and α2 secondary structure elements. This density is too large to accommodate two Mg^2+^ ions, but is well fit by a single phosphate group and is likely an artifact of the purification conditions (**Figure 1B inset**).

### PXL selectively binds G-rich ssRNA and ssDNA sequences

Building on the observation that PXL, like its homolog Exu, is a pseudo-nuclease, we next tested whether the PXL domain can directly interact with nucleic acids using electrophoretic mobility shift assays (EMSAs). In the assays, 5’-AF488-conjugated DNA and RNA oligonucleotides were incubated with increasing concentrations of PXL. EMSAs revealed that PXL has a clear preference for RNA over ssDNA of the same oligonucleotide composition (**Figure 3A-B**) and does not bind dsDNA (**Supplementary Figure S3**). Furthermore, PXL has strong RNA sequence specificity. It efficiently binds poly(UG) RNA that was previously reported as a preferred RNA substrate of Exu (32), but not poly(A), poly(C), or poly(AC), and only weakly, poly(U) RNA. Interestingly, PXL’s poly(UG) affinity further increased with the increasing number of the (UG) repeats, suggesting that there is a minimal size requirement for optimal PXL binding (**Figure 3B**). Technical limitations prevented the use of poly(G) RNA, but poly(G) ssDNA effectively associated with PXL in EMSA assays (**Figure 3A**). These data suggest that PXL can bind RNA and ssDNA *in vitro* and has a preference for G-rich motifs.

**Figure 3:**
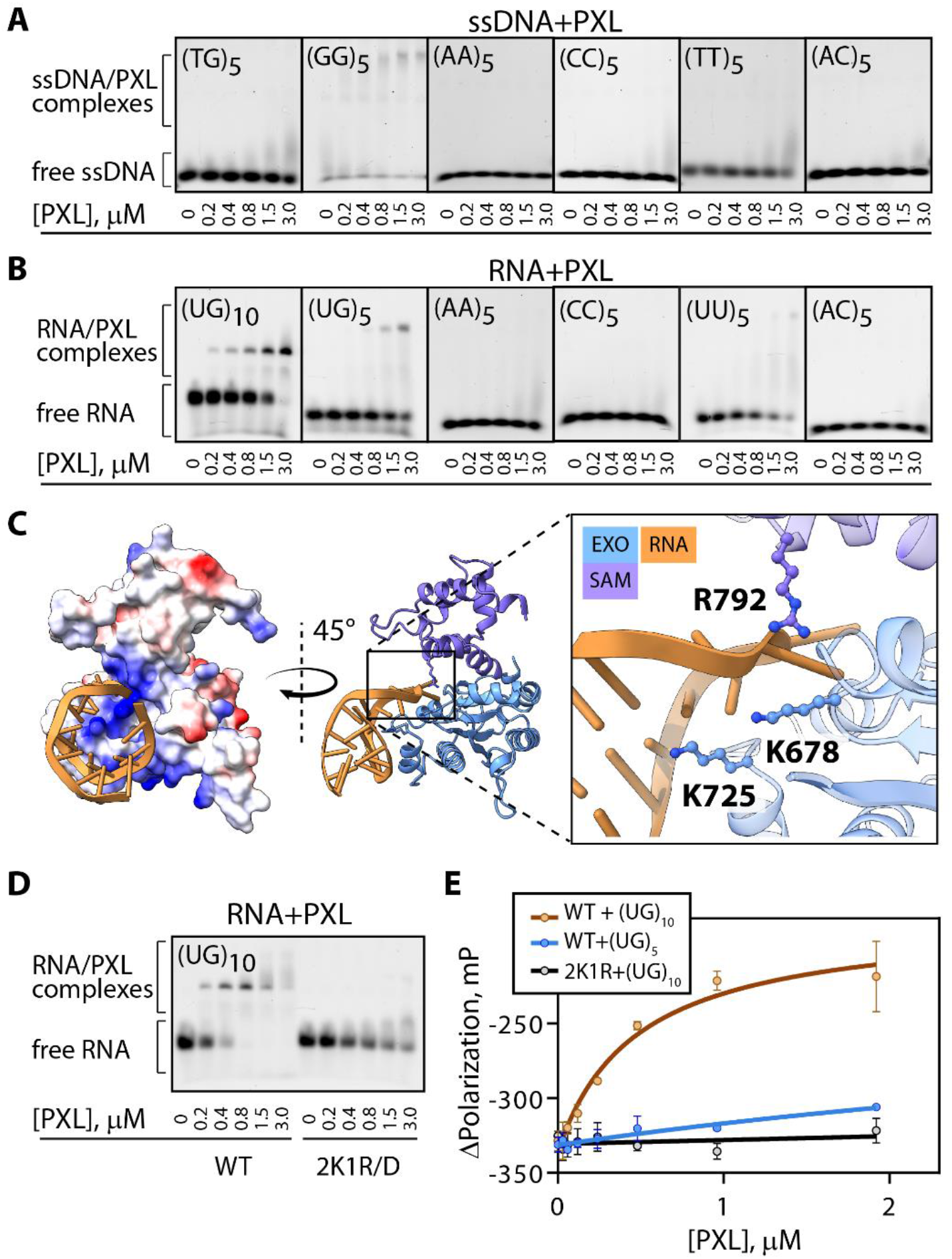
PXL binds to specific G-rich nucleic acid sequences using positively charged surface exposed residues. **(A, B)** Electrophoretic mobility shift assays of PXL and different (A)ssDNA/(B)RNA substrates. PXL demonstrates a clear preference for pUG RNA sequences and G-rich ssDNA motifs. **(C)** AlphaFold3 predicted model of PXL binding UG_(10)_ (yellow). Positively charged residues located on both the SAM-like and EXO-like subdomains used to generate PXL 2K1R/3D mutant are represented as ball and stick in the inset. **(D)** EMSA assay comparing binding of PXL and PXL 2K1R/3D to UG_(10)_ RNA. Mutation of positively charged residues on surface of PXL significantly reduces binding efficiency. **(E)** Fluorescence polarization binding curves of PXL and PXL (2K1R/3D) on poly(UG) sequences of different lengths.

The EMSA results were further validated with the fluorescence polarization binding assays (**Figure 3E** and **Supplementary Figure S3**). In these assays, PXL demonstrated robust nanomolar binding affinity for 20-nucleotide-long RNA (UG)_10_ (477±142 nM) and weaker affinity for (UG)_5_ RNA and (GG)_5_ ssDNA oligonucleotides. No binding to dsDNA was detected.

Poly(UG) RNA motifs have been reported to form stable four-stranded helical quadruplex structures, called PUG-folds, stabilized by hydrogen bonding between tetrads of guanine bases (33-37). Similar structures are formed by poly(G) repeats of ssDNA (G-quadruplexes) (38). The observed preference of PXL for G-rich motifs capable of forming these structures strongly suggests that PXL recognizes not only specific oligonucleotide sequences but may have even further specificity towards their secondary structure.

### PXL’s positively charged surface is required for its nucleic acid-binding

The PXL domain contains a positively charged surface surrounding its pseudo-active site (**Figure 1D**), making it a likely site for nucleotide binding. This hypothesis is supported by an AlphaFold model of PXL in complex with (UG)_10_ RNA (**Figure 3C**). To test whether this surface mediates RNA binding, we mutated positively charged residues that constitute the putative RNA-binding site (**Figure 3C inset**) and examined the RNA-binding ability of the resulting mutant.

Residues K678, K725, and R792, located on loops of the EXO-like domain between β4-α2 and α4-α5, as well as on α9 of the SAM-like domain, were selected based on the complex model and mutated to aspartates (PXL 2K1R/3D). An EMSA assay comparing WT PXL and the 2K1R/3D mutant showed that these mutations abolished RNA binding (**Figure 3D**). These results were further validated using FP assays, which confirmed that substrate binding was effectively eliminated by 2K1R/3D mutations (**Figure 3E**).

Together, these data demonstrate that the positively charged surface surrounding the pseudo-active site of PXL is required for its nucleic acid-binding activity.

### PXL impacts the transcriptome of U2OS cells

Given the involvement of the PXL domain in sequence-specific nucleotide binding, together with the established roles of PUG RNA motifs and G-quadruplexes in gene expression (34,38), we hypothesized that PXL may play a role in transcriptional regulation. To assess the broader impact of the PXL domain on the cellular transcriptome, we performed RNA-seq analysis.

Transcriptomes of four U2OS cell lines were analyzed and compared: WT U2OS cells expressing all PML isoforms (WT), PML knock-out cells expressing no PML (PML-KO) (26) and PML-KO cells transfected with either isoform 1 (PML-1) or isoform 1 lacking residues 601-882 (ΔPXL) (**Figure 4, Supplementary Figure S4, and Supplementary Table S1)**

**Figure 4:**
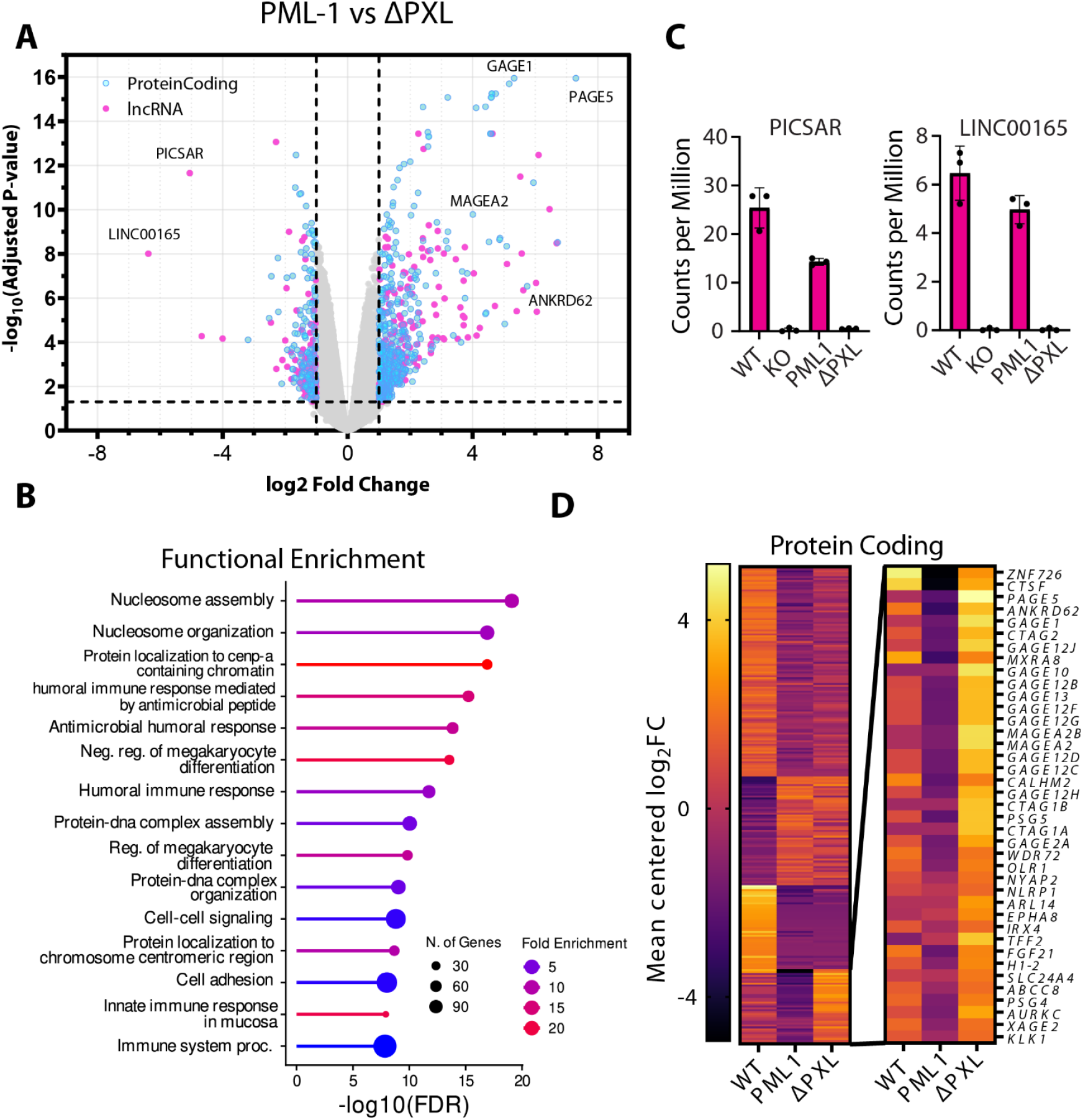
PXL affects the abundance of both protein coding and lncRNA in the U2OS transcriptome. **(A)** Volcano plot of gene expression profiles comparing U2OS cells expressing PML-1 to PML-1ΔPXL. **(B)** Abundance counts of lncRNA’s PICSAR and LINC00165 in WT U2OS, PML^-/-^, PML-1 and PML- 1ΔPXL cells. **(C)** ShinyGO enrichment analysis (39) depicting GO biological processes enriched in all differentially expressed genes between PML-1ΔPXL and PML-1. **(D)** Heatmap depicting clustering of 250 differentially expressed genes across WT U2OS, PML_-/-_, PML-1 and PML-1ΔPXL cells. Genes specifically regulated by PXL are shown in the inset.

Gene ontology enrichment analysis (39) of differentially expressed protein-coding genes between PML-1 and ΔPXL identified biological processes associated with nucleosome assembly and organization, chromatin localization, and the immune response, in good agreement with biological pathways relevant to PML and PML nuclear bodies (6,40) (**Figure 4B**).

Strikingly, lncRNAs were among the top downregulated transcripts in ΔPXL cells. For instance, the oncogenic lncRNAs PICSAR and LNC00165 (41,42) exhibited pronounced expression changes that were largely dependent on the presence of the PXL domain **(Figure 4C**). Both transcripts were expressed at high levels in WT U2OS cells and in cells expressing PML-1 but were markedly reduced in the PML KO and ΔPXL cells. These data indicate that PML-1, but not ΔPXL, can rescue the loss of PICSAR and LNC00165 expression in KO cells, and demonstrate that the PXL domain is necessary for their expression.

Many protein-coding genes were differentially expressed between PML-1 and ΔPXL cell lines, with PAGE5, GAGE1, and MAGEA2 among the most strongly upregulated following loss of the PXL domain. Importantly, several previously validated PML-binding targets were among the top hits, including MAGEA2, which has been shown to directly interact with PML and regulate its function (43), lending confidence to the newly identified targets.

To further define protein-coding transcripts regulated by the PXL domain, we performed clustering analysis of RNA abundance changes across all U2OS cell lines relative to PML-KO cells (**Figure 4D**). K-means clustering of 250 genes with the highest statistically significant fold changes across all experiments identified a subset of genes that were downregulated in PML-KO cells and rescued by PML-1 expression, but not PML-1 lacking the PXL domain (**Figure 4D inset**). These findings suggest that this group of protein-coding genes is directly or indirectly regulated by the PXL domain. Notably, approximately half of these genes belong to the members of the GAGE, PAGE, XAGE, and MAGE gene families, all of which are located on the X chromosome, suggesting a potential link between PML-1 and the X chromosome-associated transcriptional regulation (44).

Overall, these transcriptomic data demonstrate that the PXL domain has a profound impact on the cellular transcriptome and plays a key role in regulating both protein-coding genes and lncRNAs.

Taken together, the X-ray crystallography, biochemical, biophysical, and cellular assays, clearly demonstrate that PML contains a nucleic acid-binding module, which specifically binds single-stranded G-rich RNA and DNA motifs *in vitro* and influences the cellular transcriptome, which has implications for transcriptional regulation, antiviral defense, and genome maintenance mediated by PML.

## DISCUSSION

PML-1 is the most abundantly expressed PML isoform in humans and is known to play roles in the anti-viral response, nucleolar targeting, cell senescence, and the maintenance of genomic stability; however, the molecular mechanisms underlying these functions remain poorly understood. Here, we present the first crystal structure of the isoform-defining C-terminal domain of PML-1 and call it PXL. The domain has a DEDDh exonuclease structural fold but lacks the conserved active site and the nuclease activity. Instead, we demonstrate that PML can directly interact with nucleic acids and regulate the cellular transcriptome. Although PML has been previously implicated in transcriptional regulation activity through the resident proteins of PML-nuclear bodies, some of which are transcription factors(4), this work, for the first time, provides strong evidence that PML-1 itself can directly and specifically interact with nucleic acids.

We show that PXL binds nucleic acids in a sequence-specific manner, with the highest affinity for ssRNA containing UG repeats and weaker binding to ssDNA G-repeats, consistent with recognition of structured RNA. This poly(UG) specificity mirrors that of PXL’s closest structural homologue, *Drosophila* Exu, a regulator of mRNA localization and expression in early embryos (32). It is likely, in part, bestowed by the presence of its SAM-like subdomain, which is not found on most exonucleases. Residues from both the SAM-like domain and EXO-like domain of Exu contribute to RNA binding (32); similarly, R792 of PXL is located on the SAM-like domain and contains a surface-exposed arginine, which we demonstrate impacts the ability of PXL to bind nucleic acid substrates.

Poly(UG) repeats are often enriched near 5’- and 3’-splice sites of pre-mRNA and are known in C.*elegans* to form a unique G4-quadruplex structure called the pUG fold, which marks the 3’ end of mRNA as vectors for gene silencing (37). Additionally, G-repeat tetrads in DNA are known to form G-quadruplex folds (38); these quadruplexes can be intrastrand or interstrand and are biologically significant in processes such as replication, transcription, and telomere stability. Notably, G4-quadruplex DNA and RNA have been shown to colocalize with PML-NBs (45), further supporting our observations and suggesting that PXL may engage quadruplex-forming nucleic acids as part of its gene regulatory function.

In line with specificity for structured regulatory repeats and the previously described role of PML as a transcription factor, our transcriptomic data demonstrates that the PXL domain of PML-1 participates in transcriptional regulation, as there is a marked difference in transcriptomic profiles between cells expressing WT PML-1 and those expressing PML-1ΔPXL. Remarkably, increased expression of isoform 1 of PML, specifically, was recently linked to a poor prognosis in estrogen receptor-positive breast cancers, and loss of PML was shown to reduce breast cancer stem cell-related gene expression, suggesting that PML-1 can be a promising anti-cancer therapeutic target (13). In agreement with these reports, our transcriptomic data show that deletion of the PXL domain from PML-1 can decrease the abundance of PAGE/GAGE/XAGE and MAGEA transcripts, which are highly expressed in cancers (46,47). While the mechanisms by which PML regulates gene expression are likely complex, our findings indicate that the PXL domain contributes substantially to PML’s function as a transcriptional regulator and suggest that PXL itself may represent a potential target for anti-cancer therapeutic intervention.

Together, this work identifies the PXL domain as a structurally distinct nucleic acid-binding module of PML-1 that links selective recognition of structured nucleic acids to transcriptional regulation, with broad implications for understanding PML function in genome regulation, antiviral defense, and cancer biology.

## Abbreviations

PML: Promyelocytic Leukemia Protein
SUMO: small ubiquitin-like modifier
ssDNA: single-strand DNA
ssRNA: single strand RNA
ChIP: chromatin immunoprecipitation
BME: beta-mercaptoethanol
TEV: Tobacco Etch Virus
lncRNA: long noncoding RNA

## ACKNOWLEDGEMENTS

The authors wish to thank Dr. Andrea Makkay for her assistance in setting up initial EMSA assays, Dr. Evan Jellison and the UConn Health Flow Cytometry Core Facility for their assistance in creating YFP-PML cell lines, and Dr. Ann Cowan and Dr. Dennis Wright for helpful discussions. This work was supported by NIH grants R35GM156397 to IB and R21AI135451 to IB and SKW.

## CRediT author statement

I.B., S.K.W., and D.F. conceptualization; I.B., S.K.W., and D.F. methodology; D.F., I.S., D.G.B., R.S., B.H., and Y.L. investigation. I.B. and D.F. writing-original draft; I.B., S.K.W, D.F., B.H. writing-review & editing. I.B., D.F., Y.L. visualization. I.B., and S.K.W supervision; I.B. and S.K.W. project administration. I.B., and S.K.W. funding acquisition.

## Conflict of Interest None declared

## Data Availability

The expression plasmid has been deposited to and are available from the Addgene database: PML-1(578-861) (#253216). All other data is available upon request. The atomic coordinates and structure factors of PXL(578-861Δ5) have been deposited in the Protein Data Bank (http://www.wwpdb.org/) with PDB ID code 10TH. The RNA sequencing data have been deposited in the Sequence Read Archive (SRA) under accession #PRJNA1423671.

## REFERENCES

1. Chu, Y. and Yang, X. (2011) SUMO E3 ligase activity of TRIM proteins. Oncogene, 30, 1108–1116.

2. Ishov, A.M., Sotnikov, A.G., Negorev, D., Vladimirova, O.V., Neff, N., Kamitani, T., Yeh, E.T., Strauss, J.F., 3rd and Maul, G.G. (1999) PML is critical for ND10 formation and recruits the PML-interacting protein daxx to this nuclear structure when modified by SUMO-1. J Cell Biol, 147, 221–234.

3. Jan Fada, B., Reward, E. and Gu, H. (2021) The Role of ND10 Nuclear Bodies in Herpesvirus Infection: A Frenemy for the Virus? Viruses, 13.

4. Zhong, S., Salomoni, P. and Pandolfi, P.P. (2000) The transcriptional role of PML and the nuclear body. Nat Cell Biol, 2, E85–90.

5. Nisole, S., Maroui, M.A., Mascle, X.H., Aubry, M. and Chelbi-Alix, M.K. (2013) Differential Roles of PML Isoforms. Front Oncol, 3, 125.

6. Alandijany, T., Roberts, A.P.E., Conn, K.L., Loney, C., McFarlane, S., Orr, A. and Boutell, C. (2018) Distinct temporal roles for the promyelocytic leukaemia (PML) protein in the sequential regulation of intracellular host immunity to HSV-1 infection. PLoS Pathog, 14, e1006769.

7. Scherer, M., Read, C., Neusser, G., Kranz, C., Kuderna, A.K., Muller, R., Full, F., Worz, S., Reichel, A., Schilling, E.M. et al. (2022) Dual signaling via interferon and DNA damage response elicits entrapment by giant PML nuclear bodies. Elife, 11.

8. Kiesslich, A., von Mikecz, A. and Hemmerich, P. (2002) Cell cycle-dependent association of PML bodies with sites of active transcription in nuclei of mammalian cells. J Struct Biol, 140, 167–179.

9. Boutell, C., Cuchet-Lourenco, D., Vanni, E., Orr, A., Glass, M., McFarlane, S. and Everett, R.D. (2011) A viral ubiquitin ligase has substrate preferential SUMO targeted ubiquitin ligase activity that counteracts intrinsic antiviral defence. PLoS Pathog, 7, e1002245.

10. Boutell, C., Sadis, S. and Everett, R.D. (2002) Herpes simplex virus type 1 immediate-early protein ICP0 and is isolated RING finger domain act as ubiquitin E3 ligases in vitro. J Virol, 76, 841–850.

11. Cuchet-Lourenco, D., Vanni, E., Glass, M., Orr, A. and Everett, R.D. (2012) Herpes simplex virus 1 ubiquitin ligase ICP0 interacts with PML isoform I and induces its SUMO-independent degradation. J Virol, 86, 11209–11222.

12. Condemine, W., Takahashi, Y., Zhu, J., Puvion-Dutilleul, F., Guegan, S., Janin, A. and de The, H. (2006) Characterization of endogenous human promyelocytic leukemia isoforms. Cancer Res, 66, 6192–6198.

13. Pai, C.P., Wang, H., Seachrist, D.D., Agarwal, N., Adams, J.A., Liu, Z., Keri, R.A., Cao, K., Schiemann, W.P. and Kao, H.Y. (2024) The PML1-WDR5 axis regulates H3K4me3 marks and promotes stemness of estrogen receptor-positive breast cancer. Cell Death Differ, 31, 768–778.

14. Condemine, W., Takahashi, Y., Le Bras, M. and de The, H. (2007) A nucleolar targeting signal in PML-I addresses PML to nucleolar caps in stressed or senescent cells. J Cell Sci, 120, 3219–3227.

15. Mathavarajah, S., Vergunst, K.L., Habib, E.B., Williams, S.K., He, R., Maliougina, M., Park, M., Salsman, J., Roy, S., Braasch, I. et al. (2023) PML and PML-like exonucleases restrict retrotransposons in jawed vertebrates. Nucleic Acids Res, 51, 3185–3204.

16. Cuchet, D., Sykes, A., Nicolas, A., Orr, A., Murray, J., Sirma, H., Heeren, J., Bartelt, A. and Everett, R.D. (2011) PML isoforms I and II participate in PML-dependent restriction of HSV-1 replication. J Cell Sci, 124, 280–291.

17. Kabsch, W. (2010) Xds. Acta Crystallogr D Biol Crystallogr, 66, 125–132.

18. Winter, G. and McAuley, K.E. (2011) Automated data collection for macromolecular crystallography. Methods, 55, 81–93.

19. Vonrhein, C., Flensburg, C., Keller, P., Sharff, A., Smart, O., Paciorek, W., Womack, T. and Bricogne, G. (2011) Data processing and analysis with the autoPROC toolbox. Acta Crystallogr D Biol Crystallogr, 67, 293–302.

20. McCoy, A.J., Grosse-Kunstleve, R.W., Adams, P.D., Winn, M.D., Storoni, L.C. and Read, R.J. (2007) Phaser crystallographic software. J Appl Crystallogr, 40, 658–674.

21. Agirre, J., Atanasova, M., Bagdonas, H., Ballard, C.B., Basle, A., Beilsten-Edmands, J., Borges, R.J., Brown, D.G., Burgos-Marmol, J.J., Berrisford, J.M. et al. (2023) The CCP4 suite: integrative software for macromolecular crystallography. Acta Crystallogr D Struct Biol, 79, 449–461.

22. Abramson, J., Adler, J., Dunger, J., Evans, R., Green, T., Pritzel, A., Ronneberger, O., Willmore, L., Ballard, A.J., Bambrick, J. et al. (2024) Accurate structure prediction of biomolecular interactions with AlphaFold 3. Nature, 630, 493–500.

23. Murshudov, G.N., Vagin, A.A. and Dodson, E.J. (1997) Refinement of macromolecular structures by the maximum-likelihood method. Acta Crystallogr D Biol Crystallogr, 53, 240–255.

24. Williams, C.J., Headd, J.J., Moriarty, N.W., Prisant, M.G., Videau, L.L., Deis, L.N., Verma, V., Keedy, D.A., Hintze, B.J., Chen, V.B. et al. (2018) MolProbity: More and better reference data for improved all-atom structure validation. Protein Sci, 27, 293–315.

25. Pettersen, E.F., Goddard, T.D., Huang, C.C., Couch, G.S., Greenblatt, D.M., Meng, E.C. and Ferrin, T.E. (2004) UCSF Chimera--a visualization system for exploratory research and analysis. J Comput Chem, 25, 1605–1612.

26. Collin, V., Gravel, A., Kaufer, B.B. and Flamand, L. (2020) The Promyelocytic Leukemia Protein facilitates human herpesvirus 6B chromosomal integration, immediate-early 1 protein multiSUMOylation and its localization at telomeres. PLoS Pathog, 16, e1008683.

27. Erdos, G., Pajkos, M. and Dosztanyi, Z. (2021) IUPred3: prediction of protein disorder enhanced with unambiguous experimental annotation and visualization of evolutionary conservation. Nucleic Acids Res, 49, W297–W303.

28. Jumper, J., Evans, R., Pritzel, A., Green, T., Figurnov, M., Ronneberger, O., Tunyasuvunakool, K., Bates, R., Zidek, A., Potapenko, A. et al. (2021) Highly accurate protein structure prediction with AlphaFold. Nature, 596, 583–589.

29. Yang, W. (2011) Nucleases: diversity of structure, function and mechanism. Q Rev Biophys, 44, 1–93.

30. Holm, L., Laiho, A., Toronen, P. and Salgado, M. (2023) DALI shines a light on remote homologs: One hundred discoveries. Protein Sci, 32, e4519.

31. Zhou, W., Richmond-Buccola, D., Wang, Q. and Kranzusch, P.J. (2022) Structural basis of human TREX1 DNA degradation and autoimmune disease. Nat Commun, 13, 4277.

32. Lazzaretti, D., Veith, K., Kramer, K., Basquin, C., Urlaub, H., Irion, U. and Bono, F. (2016) The bicoid mRNA localization factor Exuperantia is an RNA-binding pseudonuclease. Nat Struct Mol Biol, 23, 705–713.

33. Monchaud, D. (2023) Why does the pUG tail curl? Mol Cell, 83, 330–331.

34. Petersen, R.J., Vivek, R., Tonelli, M., Roschdi, S. and Butcher, S.E. (2025) The structure, folding kinetics, and dynamics of long poly(UG) RNA. Nucleic Acids Res, 53.

35. Roschdi, S., Kume, T., Petersen, R.J., McCann, A., Escobar, C.A., Richard, A. and Butcher, S.E. (2025) Sequence and ionic requirements of pUG fold quadruplexes. bioRxiv.

36. Roschdi, S., Montemayor, E.J., Vivek, R., Bingman, C.A. and Butcher, S.E. (2024) Self-assembly and condensation of intermolecular poly(UG) RNA quadruplexes. Nucleic Acids Res, 52, 12582–12591.

37. Roschdi, S., Yan, J., Nomura, Y., Escobar, C.A., Petersen, R.J., Bingman, C.A., Tonelli, M., Vivek, R., Montemayor, E.J., Wickens, M. et al. (2022) An atypical RNA quadruplex marks RNAs as vectors for gene silencing. Nat Struct Mol Biol, 29, 1113–1121.

38. Burge, S., Parkinson, G.N., Hazel, P., Todd, A.K. and Neidle, S. (2006) Quadruplex DNA: sequence, topology and structure. Nucleic Acids Res, 34, 5402–5415.

39. Ge, S.X., Jung, D. and Yao, R. (2020) ShinyGO: a graphical gene-set enrichment tool for animals and plants. Bioinformatics, 36, 2628–2629.

40. Shastrula, P.K., Sierra, I., Deng, Z., Keeney, F., Hayden, J.E., Lieberman, P.M. and Janicki, S.M. (2019) PML is recruited to heterochromatin during S phase and represses DAXX-mediated histone H3.3 chromatin assembly. J Cell Sci, 132.

41. Luo, Y., Morgan, S.L. and Wang, K.C. (2016) PICSAR: Long Noncoding RNA in Cutaneous Squamous Cell Carcinoma. J Invest Dermatol, 136, 1541–1542.

42. Jin, Y., Wu, P., Zhao, W., Wang, X., Yang, J., Huo, X., Chen, J., De, W. and Yang, F. (2018) Long noncoding RNA LINC00165-induced by STAT3 exerts oncogenic properties via interaction with Polycomb Repressive Complex 2 to promote EMT in gastric cancer. Biochem Biophys Res Commun, 507, 223–230.

43. Peche, L.Y., Scolz, M., Ladelfa, M.F., Monte, M. and Schneider, C. (2012) MageA2 restrains cellular senescence by targeting the function of PMLIV/p53 axis at the PML-NBs. Cell Death Differ, 19, 926–936.

44. Wang, J., Shiels, C., Sasieni, P., Wu, P.J., Islam, S.A., Freemont, P.S. and Sheer, D. (2004) Promyelocytic leukemia nuclear bodies associate with transcriptionally active genomic regions. J Cell Biol, 164, 515–526.

45. Komurkova, D., Svobodova Kovarikova, A. and Bartova, E. (2021) G-Quadruplex Structures Colocalize with Transcription Factories and Nuclear Speckles Surrounded by Acetylated and Dimethylated Histones H3. Int J Mol Sci, 22.

46. Brinkmann, U., Vasmatzis, G., Lee, B. and Pastan, I. (1999) Novel genes in the PAGE and GAGE family of tumor antigens found by homology walking in the dbEST database. Cancer Res, 59, 1445–1448.

47. Kuldkepp, A., Karakai, M., Toomsoo, E., Reinsalu, O. and Kurg, R. (2019) Cancer-testis antigens MAGEA proteins are incorporated into extracellular vesicles released by cells. Oncotarget, 10, 3694–3708.

