## Supplemental Figures for "PXL: a Nucleic Acid–Binding Module of Promyelocytic Leukemia Protein"

Supplementary Information

### Supplementary figure 1

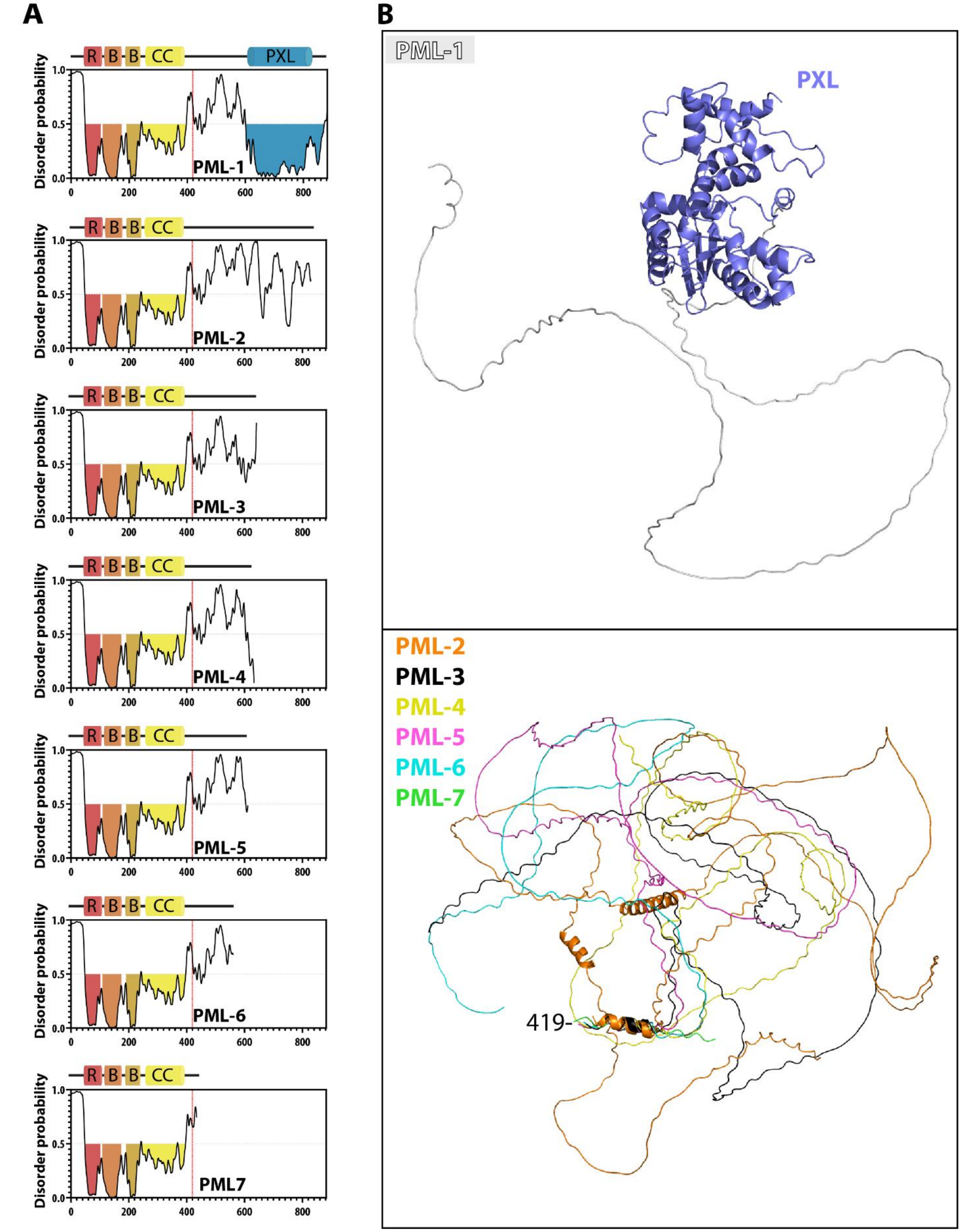


**Supplementary Figure 1: The unique C-terminus of PML-1 is predicted to contain an ordered domain.** **(A)** IUpred3 disorder predictions of all full length PML isoforms. Isoform specific C-termini begin at residue number 419 (vertical red line). PML-1 is predicted to contain an ordered region in its unique C-terminus. **(B)** AlphaFold3 predictions of PML-1 C-terminus (top) and PML-(2-7) (bottom).

### Supplementary figure 2

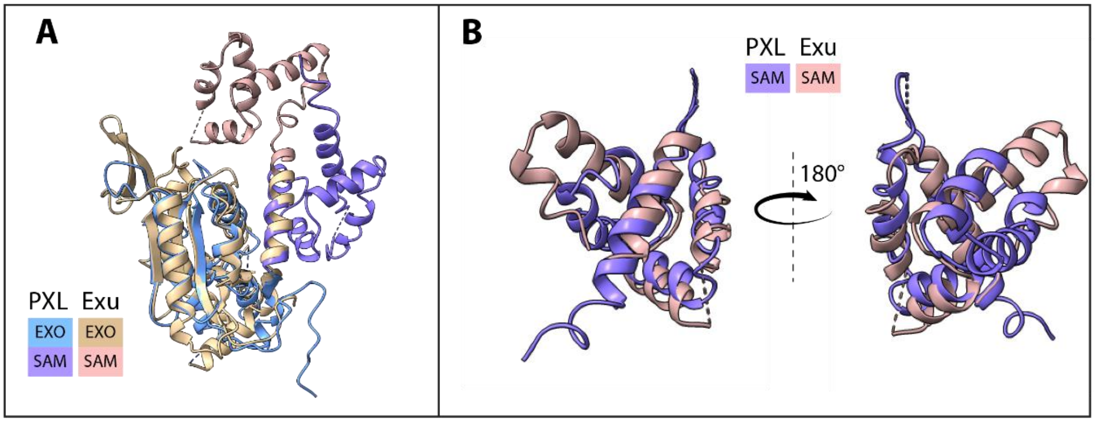


**Supplementary Figure 2: PXL and Exu SAM-like domains share secondary structure but differ in orientation relative to the Exo-like domain.**

**(A)** Overlay of the PXL and Exu exo-like (PXL: blue, Exu:Tan) and SAM-like (PXL: purple, Exu:pink) domains. The SAM-like domains differ in their orientation with respect to their Exo-like domains. **(B)** Overlay of the PXL SAM-like (purple) and Exu SAM-like (pink) domains showing conserved secondary structure alignment.

### Supplementary figure 3

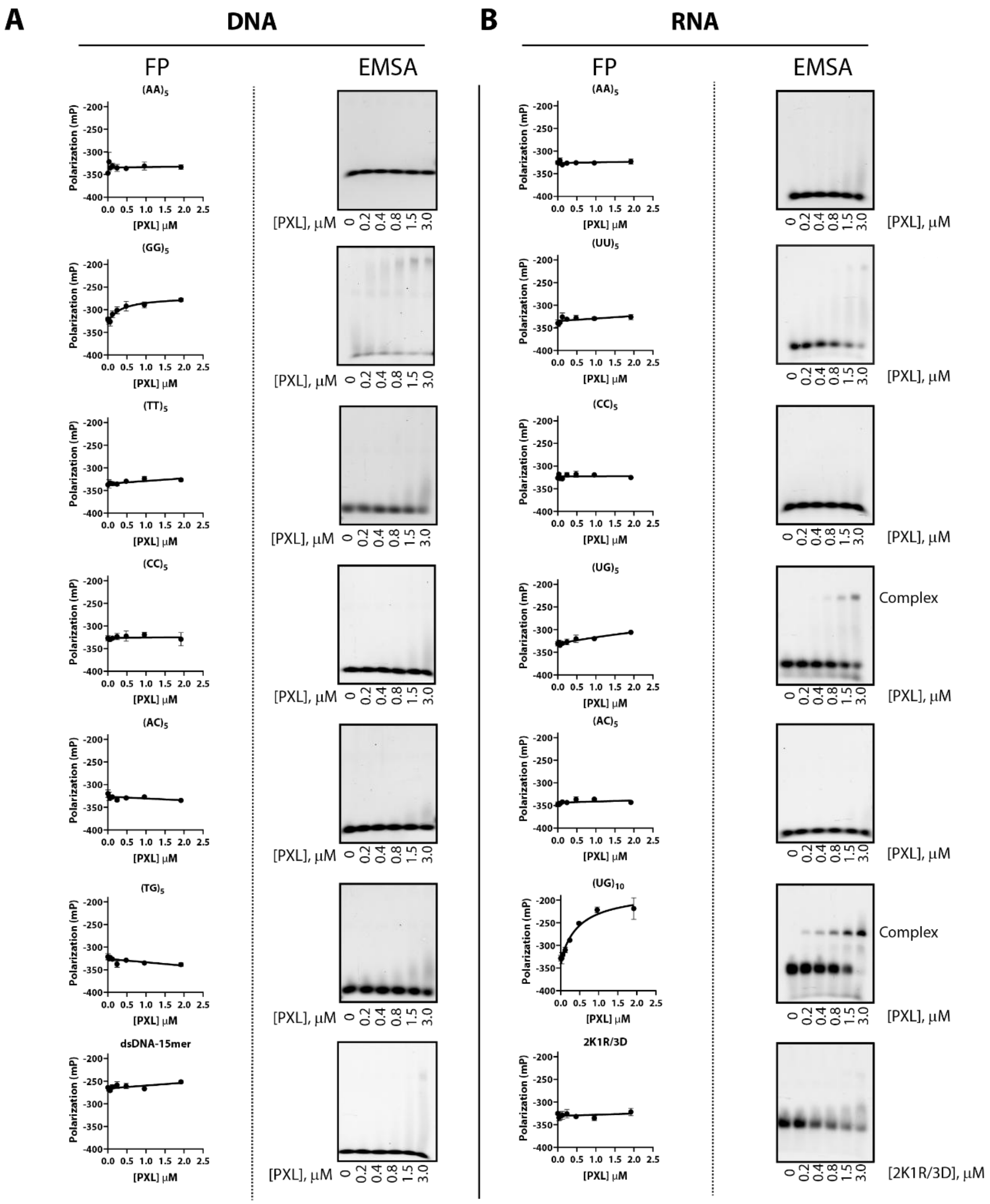


**Supplementary Figure 3: PXL selectively binds poly(UG) RNA sequences utilizing positively charged residues on both the SAM-like and Exo-like domains.**

**(A)** Fluorescence polarization (left) and electrophoretic mobility shift assays (right) of all DNA constructs used to assess PXL binding. **(B)** Fluorescence polarization (left) and electrophoretic mobility shift assays (right) of all RNA constructs used to assess PXL binding and the PXL 2K1R/3D binding deficient mutant.

### Supplementary figure 4

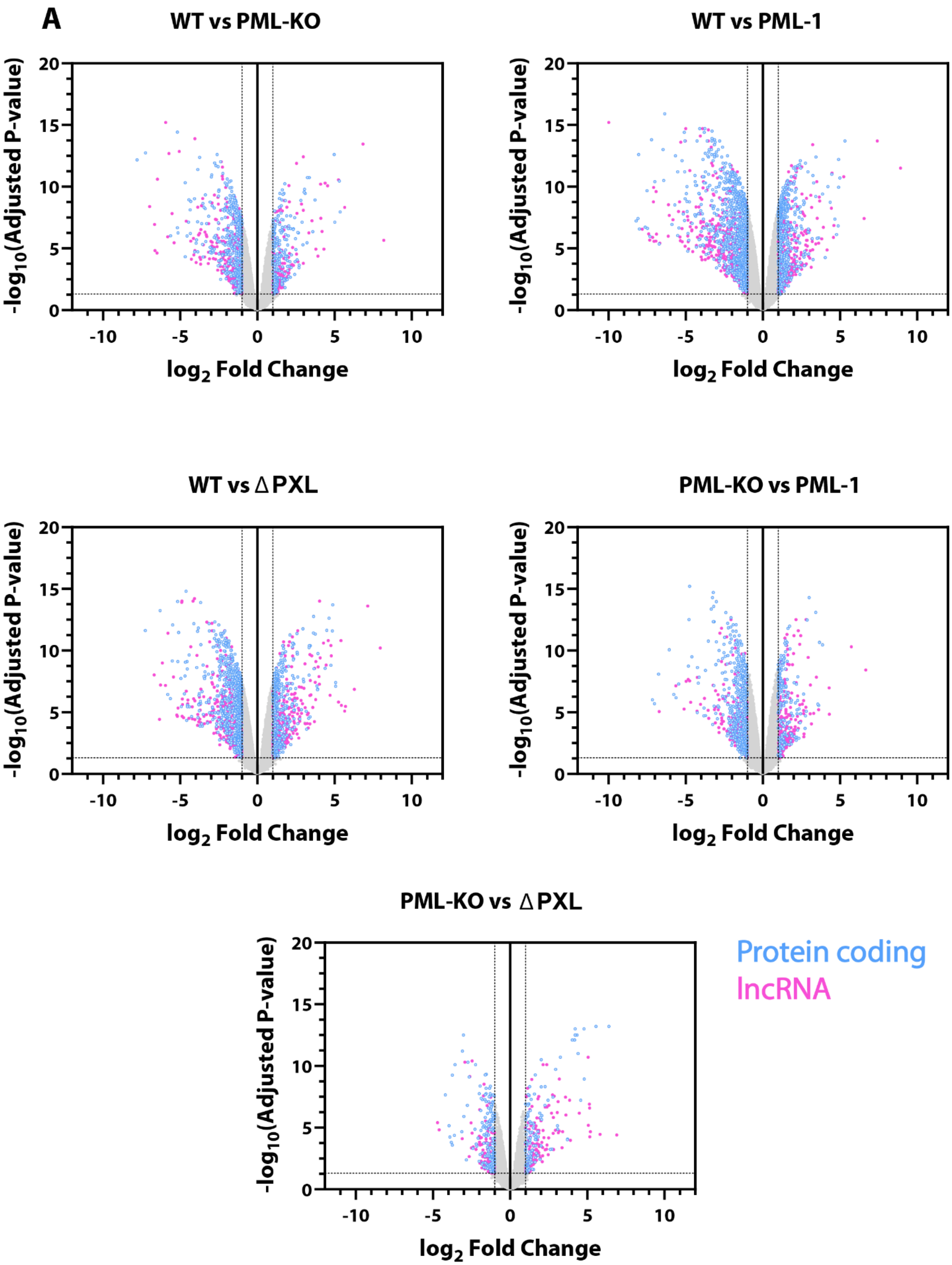


**Supplementary Figure 4: The PXL domain of PML-1 affects the transcriptome of U2OS cells.**

**(A)** Volcano plots comparing changes in transcriptomes across U2OS cell lines showing significant changes in the abundance of both lncRNAs (pink) and protein coding genes (blue).
